# Systemic and local immune response to intraocular AAV vector administration in non-human primates

**DOI:** 10.1101/2021.09.13.460058

**Authors:** Divya Ail, Duohao Ren, Elena Brazhnikova, Céline Nouvel-Jaillard, Stephane Bertin, Sylvain Fisson, Deniz Dalkara

## Abstract

The positive clinical outcomes in adeno-associated virus (AAV)-mediated retinal gene therapy have often been attributed to the low immunogenicity of AAVs along with the immune-privilege of the eye. However, several recent preclinical studies and clinical trials have shown potential for inflammatory responses to AAV mediated gene therapy. Our current understanding of the factors contributing to intraocular inflammation such as the existence of serum antibodies against AAVs prior to injection and their contribution to increases in antibody levels post-injection is incomplete. The parameters that regulate the generation of new antibodies in response to the AAV capsid or transgene post-injection after intraocular administration are also insufficiently described. In this study we carried out a retrospective analysis of the pre-existing serum antibodies in correlation with changes in antibody levels after intraocular injections of AAV in non-human primates (NHPs). We analyzed NHP serums for the presence of both Binding Antibodies (BABs), as well as a subset of these called Neutralizing Antibodies (NABs) that impede AAV transduction upon binding. We observed significantly higher pre-existing serum BABs against AAV8 compared to other serotypes. We observed a dose-dependent increase in both BABs and NABs in the serums collected post-injection, irrespective of the serotype or the mode of injection. Lastly, we were able to demonstrate a co-relation between the serum BAB levels with clinical grading of inflammation and levels of transgene expression.

## INTRODUCTION

Adeno-associated virus (AAV) is a non-enveloped virus with a single-stranded DNA genome^1^. There are 12 naturally occurring serotypes, each differing in the structure of the capsid which, in turn affects their tropism. AAV1, 4, 7, 8 and 9 have a non-human primate (NHP) origin, whereas AAV2, 3 and 5 have a human origin^2^. In addition to these, several novel AAV variants are being discovered^3^ or generated to fulfill specific needs^4,5^. Both naturally occurring and generated AAVs show distinct transduction profiles that are cell, tissue and even species specific^6^.

The first approved gene therapy consisted of the *Rpe65* gene packaged in AAV2 vector that was delivered by subretinal injections in patients with Leber’s Congenital Amaurosis (LCA)^7,8^. Following this success there have been several completed or ongoing clinical trials for AAV based gene therapy, particularly for eye diseases. A major factor that makes the eye an attractive target tissue is its relative immune-privilege, which is attributed to the presence of a blood-retina barrier, presence of local anti-inflammatory agents and myeloid cells actively counteracting adaptive immunity^9^. Although the initial reports from the clinical studies showed therapeutic efficacy, vision improvement and an excellent safety profile with the AAV, follow-up studies revealed inflammatory reactions both in preclinical studies and clinical trials^10^. Inflammatory responses can be problematic for several reasons, one of them being the potential clearance of the transduced cells by immune mechanisms bringing into question both the immune privilege and the low immunogenicity of AAV^7,10^.

In humans AAV exposure is not associated with any pathology and antibodies against AAVs are prevalent in human populations across the globe. This poses two major issues with respect to their usage for gene therapy. If a patient already has high levels of antibodies against AAVs, then injections with AAVs might trigger a stronger immune response that can potentially contribute to inflammation. Inflammation can result in the clearance of the transduced cells by the immune system that will not just reduce the efficacy of the therapy but could also worsen the condition. The second issue pertains to a subset of these total antibodies called neutralizing antibodies (NABs), which recognize and bind to the virus and neutralize it, preventing transduction, transgene expression and thus reducing efficacy^11^. A study analyzing the prevalence of different AAV serotypes in human population worldwide revealed that NABs were present against all the tested serotypes, with the prevalence against AAV1 (67%) and AAV2 (72%) being significantly higher than AAV5 (40%), AAV8 (38%) and AAV9 (47%)^12^. Another study analyzing over 800 patient samples from 4 continents and 10 countries also concluded that the NABs against AAV1 and AAV2 were higher than anti-AAV7 or anti-AAV8 NABs in humans^13^.

Ocular gene delivery can be done by intravitreal injections as well as injections in the subretinal space, which is believed to be the less immunogenic mode of injection^14,15^. However, primate studies have shown ocular immune responses following both subretinal and intravitreal modes of injection^16,17^. Other factors that can influence the immune responses are AAV serotype^18^, virus dose injected^19^, re-administration of injection^20^, promoter^21,22^ and transgene^23^.

As part of several completed and ongoing studies conducted on NHPs that involved ocular injections (both subretinal and intravitreal), we collected blood serum samples before and after the ocular AAV injections. In this study, we performed a retrospective analysis of these serum samples from animals we grouped together based on the different AAV serotypes, doses, mode of injection, and promoters used. These NHPs were also monitored for clinical signs of inflammation by slit lamp examination and fundoscopy. Eye fundus imaging and optical coherence tomography (OCT) was done to test transgene expression as well as the structural integrity of the retina and co-related to the serum antibody levels. With this retrospective analysis we confirm in NHPs some trends observed in small animal studies as well as provide new insights that can prove valuable for designing future clinical and large animal studies.

## RESULTS

### Seroprevalence of antibodies against frequently used AAV serotypes in NHP population

Studies analyzing human serums have shown a prevalence of anti-AAV1 and anti-AAV2 antibodies at a higher level than other serotypes^12,13^. We tested the basal levels of antibodies in 41 NHPs against different serotypes that are commonly used for gene delivery to the retina – AAV2, 5, 8 and 9 by ELISA. We found that in the NHP serums the level of anti-AAV8 antibodies is significantly higher than all other serotypes tested. The level of anti-AAV9 antibodies was also significantly higher than AAV2 and AAV5 but lower than AA8 (Figure 1). There was considerable variability among the individual macaques in the anti-AAV8 and anti-AAV9 groups, but the intergroup differences were significant (Supplementary figure 1). It is worth noting that a majority of the 41 animals received intraocular injections with AAV2 based vectors, and hence were sometimes pre-selected to have low levels of anti-AAV2 serum antibodies. Hence, differences we observe between anti-AAV2 antibodies and other serotypes are likely skewed. Nonetheless, none of these animals were pre-selected for low antibodies against AAV5 or AAV8, so the difference between these two serotypes reflects the seroprevalence of those serotypes in the Mauritius macaque population.

**Figure 1.**
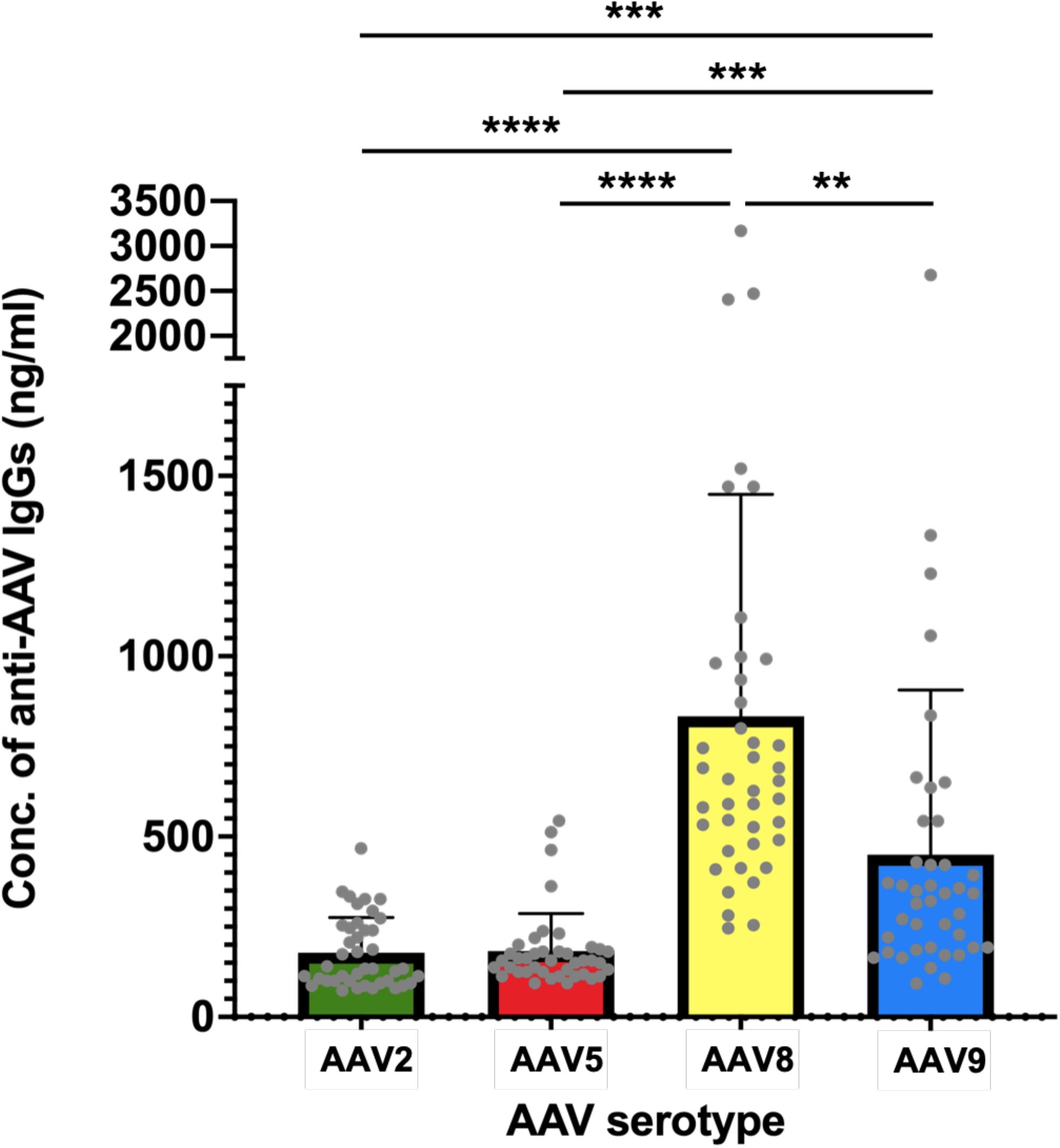
Higher basal levels of BABs against AAV8 and AAV9 in NHP serums. Mean ±SD of the concentration of BABs against AAV2 (green), AAV5 (red), AAV8 (yellow) and AAV9 (blue) in serums from 41 NHPs before injection. Significance between individual serotypes was tested using Student’s t-test (*P< 0.05, ** P< 0.01, ***P< 0.001, **** P< 0.0001).

### Increase in both the BAB and NAB levels post-injection

Next, we wanted to test the change in the anti-AAV BABs and NABs post-injection. Towards this goal, we collected the blood sample before ocular injections (BI) and post injection (PI), isolated and stored the serum component of the blood for further analysis. At each time point these serum samples were tested for BABs by ELISA and for NABs by a cell based NAB assay (Figure 2A). The NHPs received bilateral injections, and the serum samples were grouped according to the total dose received. The animals in the high dose (1-6 × 10^12^ vg) group had a 3.7-fold increase in serum BAB levels post-injection, whereas the increase was 1.8-fold in case of the animals that received a medium dose (1-6 × 10^11^ vg), and there was no significant difference in the animals that received the low dose (1-6 × 10^10^ vg) (Figure 2B). This trend was observed irrespective of the serotype injected, as well as when a combination of serotypes was injected (Supplementary figure 2). To test the NABs, we need a cell line that is effectively transduced by the different serotypes. We are able to perform NAB assays for AAV2 using HEK293T cells and for AAV9 using Lec2 cells^24^. In our cohort of animals, a majority were injected with AAV2 or AAV2-7m8, a variant of AAV2, so we grouped these by the dose of AAV2 they received and tested the levels of NABs in the serum before and post-injection. In the high dose group, there was an increase in the level of NABs post injection in all the animals tested, with 3 out of 19 animals showing very high levels which were able to neutralize the AAV even at a 5000-fold dilution (Figure C), and 16 animals with an increase comparable to the positive control (Figure D). The BAB levels in these groups showed a 5.6-and 3.5-fold increase respectively (Figure C’ and D’). Even in the group receiving the medium dose, two types of responses were observed – 3 animals showed an increase in NABs comparable to the positive control (Figure E), whereas 4 animals did not seem to have developed anti-AAV2 NABs post-injection (Figure 2F). The level of BABs in this group also co-relates to the NAB level showing a 2.5-fold increase in the medium dose-response type 1(Figure 2E’), and no change in the BAB levels in the medium dose-response type 2 group (Figure 2F’). 2 animals were injected with a low dose of AAV2, and both do not have an increase in NABs or BABs (Figure 2G and G’). The NAB level in serum is defined by the serum dilution at which 50% of the cells are neutralized. This value could not be determined for the 2 samples at low dose and the 3 samples at high dose with response type 1, so these were set at the limits of our experiment at 1/5 and 1/5000 respectively. For the high dose-response type 2, the medium dose-response type 1 and medium-dose type 2 response, this value was between 1/100 and 1/500, more precisely at 0.0085, 0.01 and 0.19 respectively (Figure 2H).

**Figure 2.**
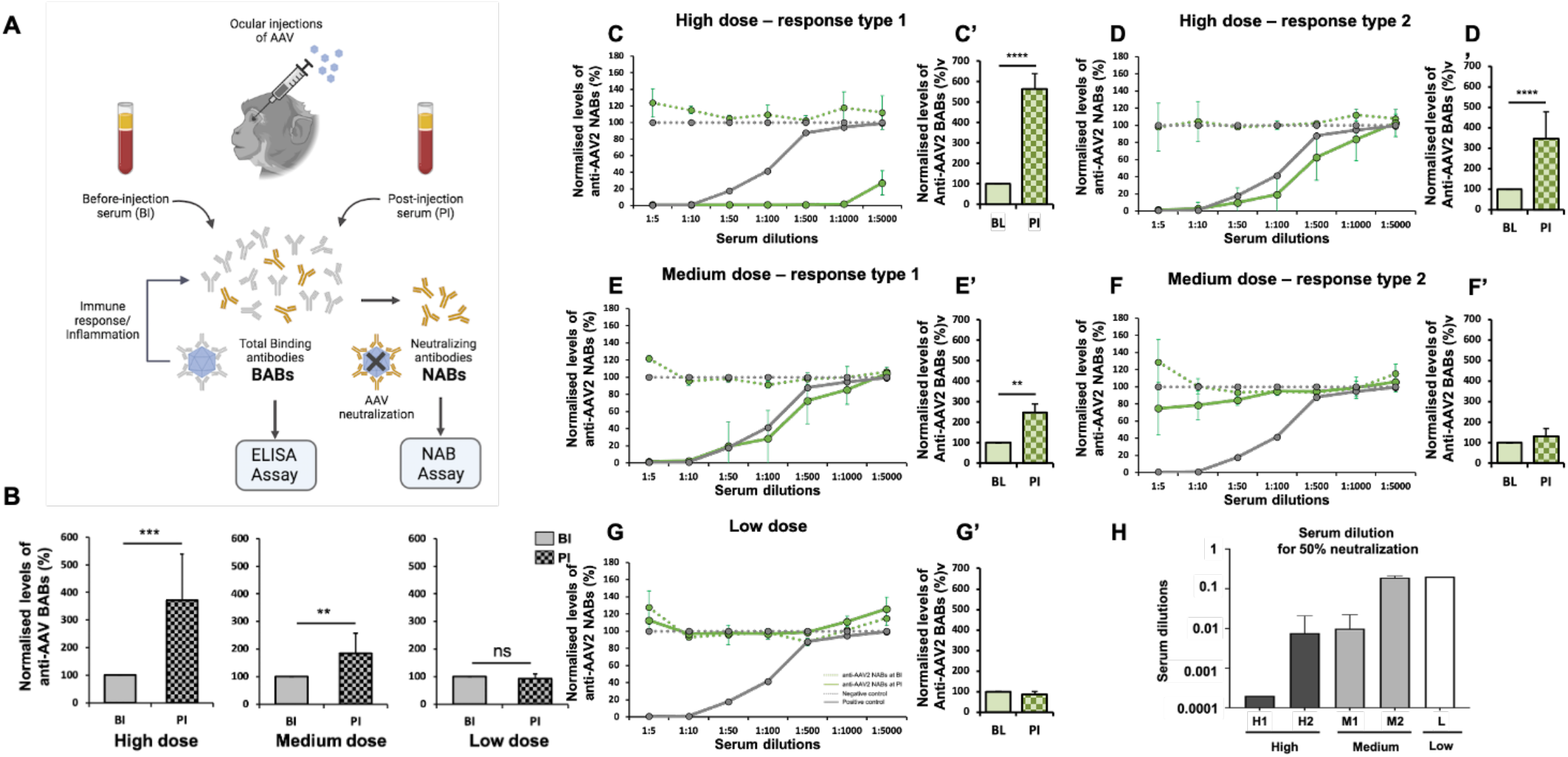
Dose dependent increase in levels of BABs and NABs post-injection. (A) Schematic representation of the experimental protocol showing the serum collection points BI (before injection) and PI (post-injection) from which the total binding antibodies (BABs) are isolated for testing by ELISA, and a subset of these, the neutralizing antibodies (NABs) are tested by NAB Assay; (B) Mean fold change in the concentration of BABs against AAV in n=26 NHPs at high dose, n=9 NHPs at medium dose and n=4 NHPs at low dose. (C-G) Change in the levels of anti-AAV2 NAB levels and (C’-G’) BAB levels in (C,C’) n=16 NHPs, (D,D’) n=3 NHPs, (E,E’) n=3 NHPs, (F,F’) n=4 NHPs and (G,G’) n=2 NHPs. (H) Serum dilution at which 50% of the AAVs are neutralized (H1 and H2: NHPs with High dose response type 1 and 2 respectively, M1 and M2: NHPs with Medium dose response type 1 and 2 respectively, L: NHPs with low dose. The values for BABs are normalized relative to the BI level (set to 100), and shown as Mean ±SD. The values for NABs at each dilution are normalized relative to the negative control (set to 100) and shown as Mean ±SD. Significance between individual time-points was tested using Student’s t-test (*P< 0.05, ** P< 0.01, ***P< 0.001, **** P< 0.0001). High dose: 10-60 × 10^11^ vg, Medium dose: 1-6 × 10^11^ vg, Low dose: 0.1-0.6 × 10^11^ vg.

### Local signs of inflammation correlate to serum antibody levels

The dose also has an impact on the retina, as deposits and structural abnormalities were observed in some of the animal eyes that received a high dose, whereas these signs did not appear in animals that received a medium or low dose. In the left eye of NHP39 that was injected with a high dose (1.00E+12vg) structural changes were observed one month post injection which persisted until 5 months (arrows pointing at the anomalies in Figure 3B’-C’). These anomalies are not observed in NHP27 that received a medium dose (5.00E+11vg). The NHP 27 and 39 were injected intravitreally with AAV2 capsid containing the same promoter and transgene (Figure 3A-F’). Further, clinical signs of local immune response were graded using the Standardization of Uveitis Nomenclature (SUN) working group and the NIH grading scales^25^. We counted the anterior chamber/ vitreous cells and evaluated signs of posterior uveitis. A summary of 4 eyes – left and right eye (from 2 NHP in each case injected with AAV2) for each dose is shown (Figure 3G). In NHP24 and NHP39 that received the high dose (1.00E+12vg) there were cells in the anterior chamber that subsided eventually, but the cells in the vitreous persisted until 5 months, albeit at a low level. Opacity of the eye or vitreal haze were only observed in eyes that received the high dose or in case of NHP25 and NHP27 that had received a medium dose (5.00E+11vg), but not in the eyes of NHP34 or NHP41, that received low dose (5.50E+10vg) injections. Posterior uveitis at a very low level of grade 1 was observed in many of the animals tested (Figure 3G).

**Figure 3.**
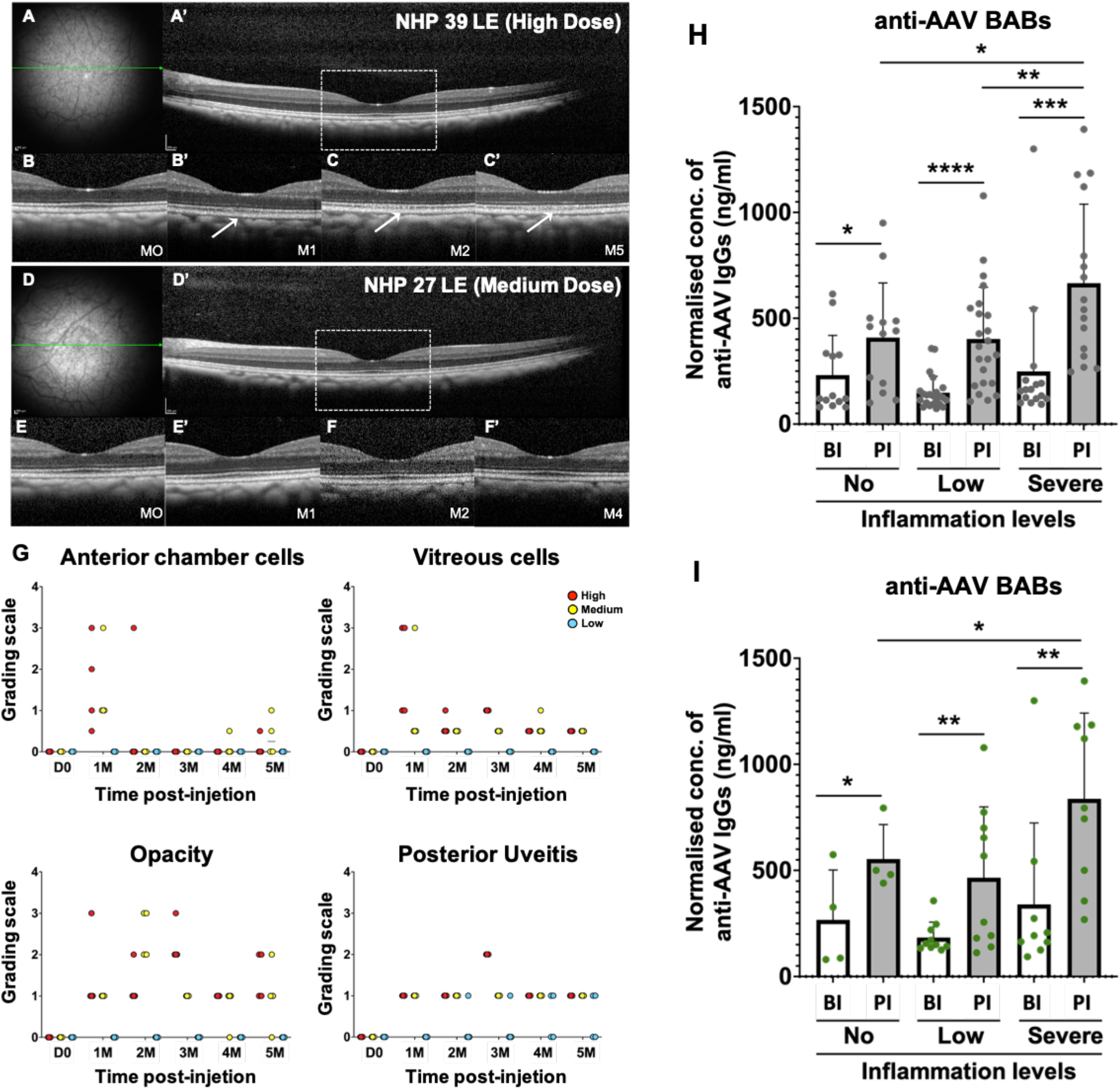
increase in the serum levels of BABs co-relates with clinical signs of inflammation. OCT images of the retina of (A-C’) NHP 39 that received a high dose injection and (D-F’) NHP 27 that received a medium dose injection; (A,D) Fundus image of the injected area; (A’,D’) Cross-section of the retina close to the fovea, marked by the green line in (A) and (D); the region demarcated by the box is shown at (B,E) the day of injection: M0, (B’,E’) 1 month post-injection:M1, (C,F) 2 months post-injection:M2 and (C’,F’) 4-5 months post-injection, as indicated; (G) Clinical grading of ocular immune response by evaluation of anterior chamber cells, vitreous cells, opacity and posterior uveitis from the day of injection (D0) to 5 months post-injection (5M); (H) Serum levels of anti-AAV BABs in NHPs before (BI) and post injection (PI), grouped by the level of ocular inflammation observed; (I) Serum levels of anti-AAV BABs in NHPs, grouped by the level of ocular inflammation and transgene expression (NHPs from Table1). The values for BABs are shown as Mean ±SD. Significance between individual time-points was tested using Student’s t-test (*P< 0.05, ** P< 0.01, ***P< 0.001, **** P< 0.0001).

Based on this clinical grading for local ocular immune responses, as well as monitoring of the health of the retina by imaging and retinal structure by optical coherence tomography (OCT), we divided the animals into 3 groups – the first showed no signs of inflammation, the second group showed some signs of inflammation and the third group included animals that we described as having severe inflammation. The animals in the first group required no intervention, the ones with low inflammation were given local kenacort injections 2-5 days post AAV injections; and animals that had severe inflammation were given additional kenacort treatment and in some cases intramuscular injections of short-term systemic corticosteroids. In each group there was a significant increase in the level of BABs post-injection compared to before injection – 1.8-fold for the no inflammation group, 2.4-fold for the low inflammation group and 2.7-fold increase for the severe-inflammation group. Interestingly there was a 1.6-fold difference between the post-injection levels of the no-inflammation and severe-inflammation group, which was significant (Figure 3H). Some of the animals in these groups were injected with a transgene that contained a GFP reporter. Hence, for these animals we looked at the transgene expression by imaging the fundus (Table 1). Some animals (NHP6, NHP8 and NHP11) in the severe-inflammation group did not express the transgene, whereas the animals that received the same serotype of AAV at the same dose but had a low level of inflammation showed good reporter gene expression (NHP6, NHP7 and NHP10). The level of BABs in these animals showed a trend like the one observed in all the animals combined (Figure 3I).

**Table 1.**
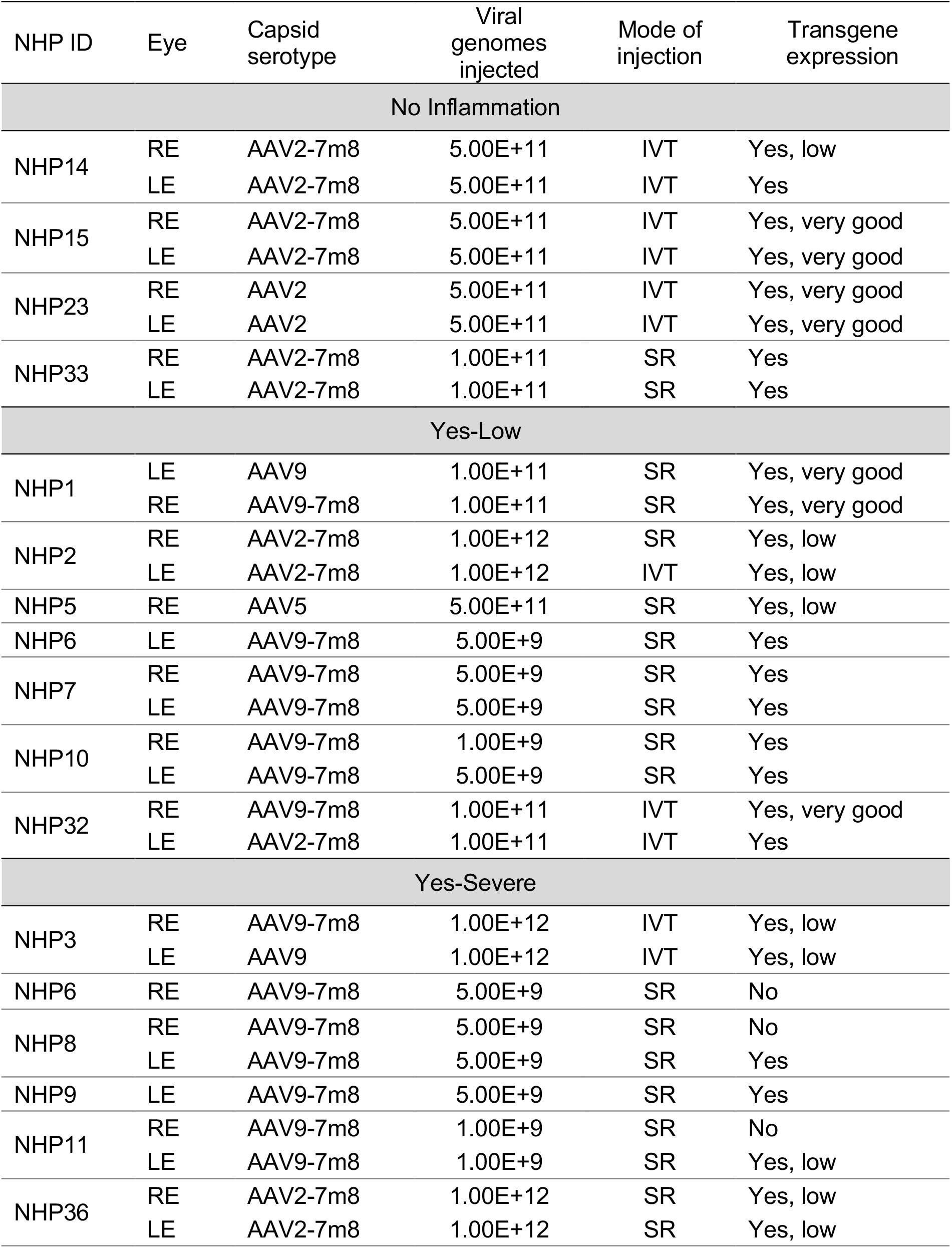
Effect of local signs of inflammation on transgene expression.

| NHP ID | Eye | Capsid serotype | Viral genomes injected | Mode of injection | Transgene expression |
| --- | --- | --- | --- | --- | --- |
| No Inflammation |  |  |  |  |  |
| NHP14 | RE | AAV2-7m8 | 5.00E+11 | IVT | Yes, low |
|  | LE | AAV2-7m8 | 5.00E+11 | IVT | Yes |
| NHP15 | RE | AAV2-7m8 | 5.00E+11 | IVT | Yes, very good |
|  | LE | AAV2-7m8 | 5.00E+11 | IVT | Yes, very good |
| NHP23 | RE | AAV2 | 5.00E+11 | IVT | Yes, very good |
|  | LE | AAV2 | 5.00E+11 | IVT | Yes, very good |
| NHP33 | RE | AAV2-7m8 | 1.00E+11 | SR | Yes |
|  | LE | AAV2-7m8 | 1.00E+11 | SR | Yes |
| Yes-Low |  |  |  |  |  |
| NHP1 | LE | AAV9 | 1.00E+11 | SR | Yes, very good |
|  | RE | AAV9-7m8 | 1.00E+11 | SR | Yes, very good |
| NHP2 | RE | AAV2-7m8 | 1.00E+12 | SR | Yes, low |
|  | LE | AAV2-7m8 | 1.00E+12 | IVT | Yes, low |
| NHP5 | RE | AAV5 | 5.00E+11 | SR | Yes, low |
| NHP6 | LE | AAV9-7m8 | 5.00E+9 | SR | Yes |
| NHP7 | RE | AAV9-7m8 | 5.00E+9 | SR | Yes |
|  | LE | AAV9-7m8 | 5.00E+9 | SR | Yes |
| NHP10 | RE | AAV9-7m8 | 1.00E+9 | SR | Yes |
|  | LE | AAV9-7m8 | 5.00E+9 | SR | Yes |
| NHP32 | RE | AAV9-7m8 | 1.00E+11 | IVT | Yes, very good |
|  | LE | AAV2-7m8 | 1.00E+11 | IVT | Yes |
| Yes-Severe |  |  |  |  |  |
| NHP3 | RE | AAV9-7m8 | 1.00E+12 | IVT | Yes, low |
|  | LE | AAV9 | 1.00E+12 | IVT | Yes, low |
| NHP6 | RE | AAV9-7m8 | 5.00E+9 | SR | No |
| NHP8 | RE | AAV9-7m8 | 5.00E+9 | SR | No |
|  | LE | AAV9-7m8 | 5.00E+9 | SR | Yes |
| NHP9 | LE | AAV9-7m8 | 5.00E+9 | SR | Yes |
| NHP11 | RE | AAV9-7m8 | 1.00E+9 | SR | No |
|  | LE | AAV9-7m8 | 1.00E+9 | SR | Yes, low |
| NHP36 | RE | AAV2-7m8 | 1.00E+12 | SR | Yes, low |
|  | LE | AAV2-7m8 | 1.00E+12 | SR | Yes, low |

### The mode of injection and its impact on the increase of anti-AAV antibodies in blood sera

The vitreous is composed of a gelatinous substance composed of water and a network of collagen and hyaluronan^11^. It has been shown that intravitreal injection results in increased spreading of AAV particles into systemic circulation leading to adaptive responses^15^. We tested the difference in BAB and NAB levels in animals that received AAV by different modes of intraocular injection. We observed a 3.6-fold increase in the level of BABs post-injection in the 6 animals that received bilateral subretinal injections, a 4.1-fold increase in 12 animals that received intravitreal injections and a 4.4-fold increase in animals that received a combination of subretinal and intravitreal injections (Figure4C). Some of these animals were injected with AAV2 and hence tested for the NAB levels prior to injection. In each group we observed that post AAV injection there was an increase in NABs which was comparable to the positive control (Figure 4B, D and F), and there was no difference in the serum dilution at which 50% neutralization is obtained (Figure 4E).

**Figure 4.**
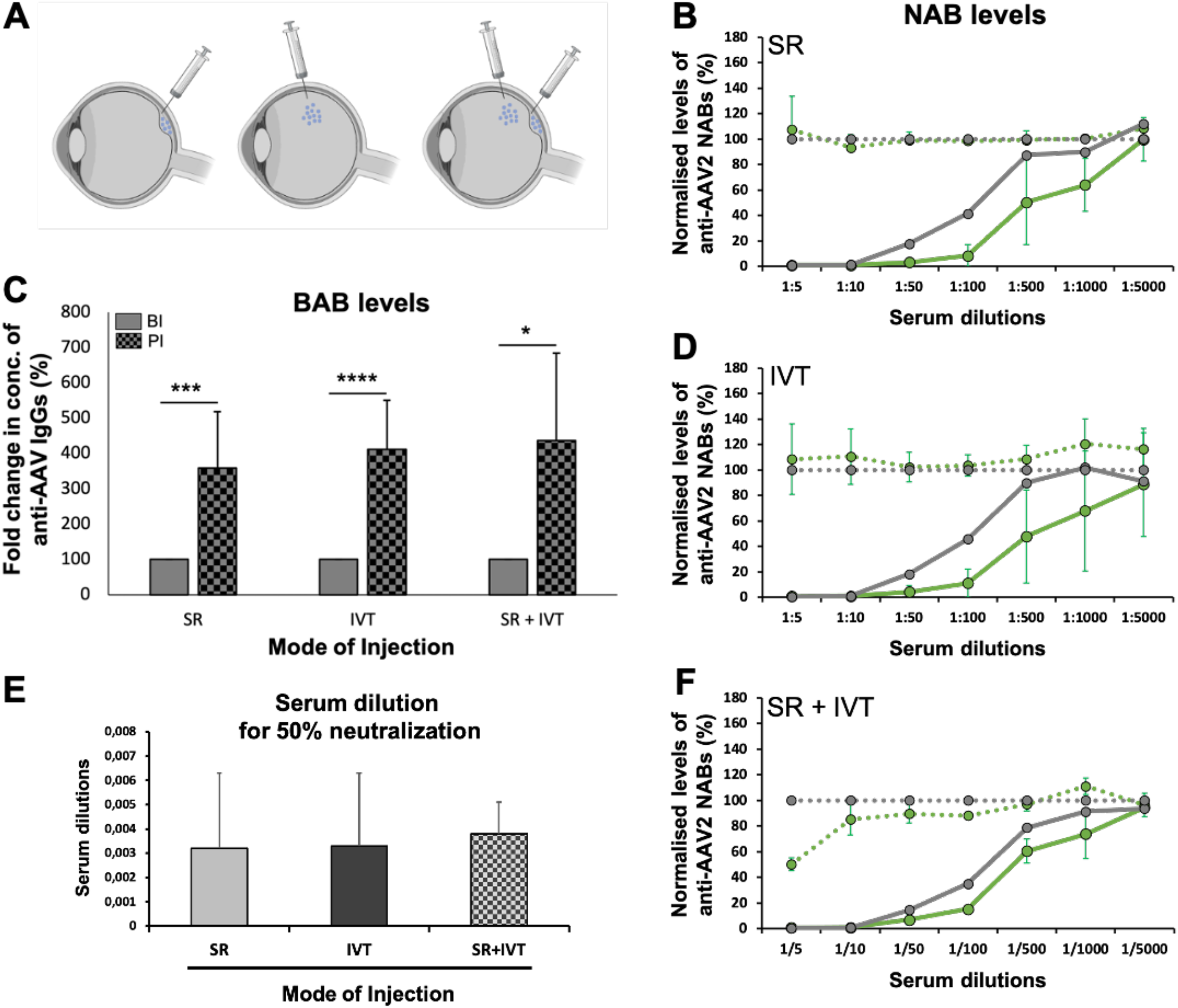
No impact of the mode of ocular injection on the serum levels of BABs and NABs. (A) Schematic representation of the modes of injection, SR: subretinal, IVT: Intravitreal; Anti-AAV2 NAB levels in (B) n=2 NHPs that received SR injections, (D) n=12 NHPs that received IVT injections and (F) n=2 NHPs that received a combination of SR and IVT injections. (C) BAB levels in n=6 NHPs that received SR injections, n=12 animals that received IVT injections and n=4 NHPs that received SR+IVT injections; (E) Serum dilution at which 50% of the AAVs are neutralized. The values for NABs are normalized relative to the negative control (set to 100), and shown as Mean ±SD. The values for BABs are normalized relative to the BI level (set to 100) and shown as Mean ±SD. Significance between individual time-points was tested using Student’s t-test (*P< 0.05, ** P< 0.01, ***P< 0.001, **** P< 0.0001).

### Pre-existing antibodies and cross-reactivity across serotypes affect antibody levels post-injection

An important consideration for injections is the pre-existing levels of BABs and NABs^26,27^. We grouped the animals based on the presence of pre-existing antibodies, and then compared the post-injection increase in BABs to the levels before injection. As expected, there was an increase post-injection which was slightly higher in the group with pre-existing BABs. But there was no significant difference between the post-injection levels between the group with and without pre-existing BABs (Figure 5A-C). The NAB and BAB levels against AAV2 increased in both NHP14 (without pre-existing NABs) and NHP36 (with pre-existing NABs). Similarly, the NAB and BAB levels against AAV9 increased in both NHP9 and NHP8 (Figure5D-G).

**Figure 5.**
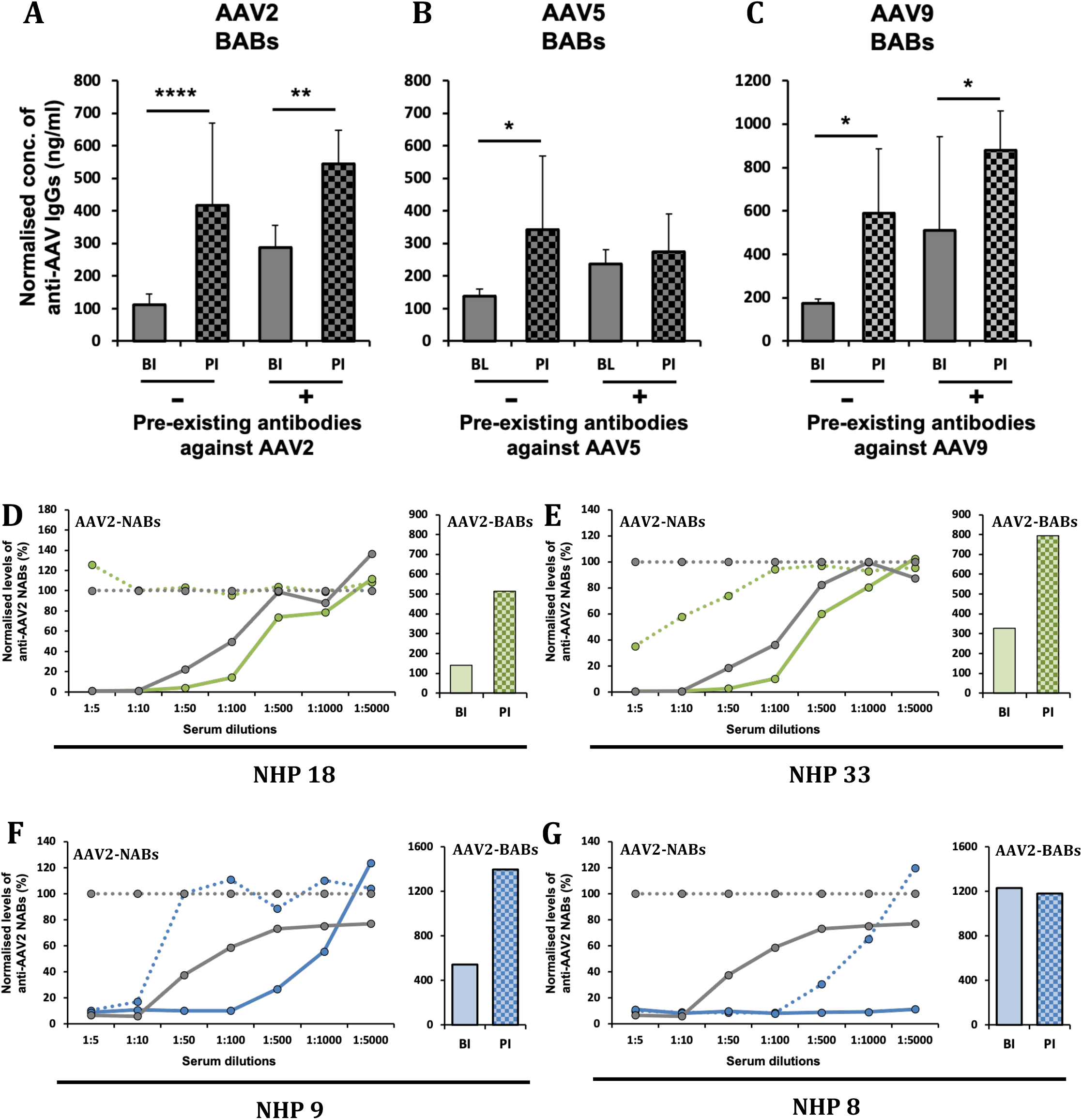
Effect of pre-existing antibodies on BAB and NAB production. Serum concentration of BABs against (A) AAV2 in n=22 NHPs without (-) and n=5 NHPs with pre-existing (+) BABs; (B) against AAV5 in n=7 NHPs without (-) and n=2 NHPs with pre-existing (+) BABs,; (C) against AAV9 in n=4 NHPs without (-) and n=7 NHPs with pre-existing (+) BABs; Serum concentration of NABs and BABs against AAV2 in (D) NHP14 and (E) NHP36; Serum concentration of NABs and BABs against AAV9 in (F) NHP9 and (E) NHP8. In each case the BABs and NABs are tested against the same serotype that the animals were injected with. The values for BABs are shown as Mean ±SD. The values for NABs at each dilution are normalized relative to the negative control (set to 100) and shown as Mean ±SD. Significance between individual time-points was tested using Student’s t-test (*P< 0.05, ** P< 0.01, ***P< 0.001, **** P< 0.0001).

Some studies have shown that there is cross-reactivity across serotypes, wherein an injection with one serotype resulted in an increase in NABs against another serotype^26,28^. To test this, we grouped the animals that had received AAV2 injections based on their pre-injection levels of BABs against AAV5, AAV8 and AAV9. We observed a significant increase in anti-AAV9 BABs in the group of 20 animals that had pre-existing anti-AAV9 BABs but were injected with AAV2 (Figure 6A-C). NHP8 had low pre-existing anti-AAV2 NAB and BAB levels, and this level remained low post-injection of a combination of AAV5 and AAV9. On the other hand, NHP9 had some pre-existing anti-AAV2 NABs and BABs that further increased post-injection with AAV5 and AAV9 (Figure 6F-G).

**Figure 6.**
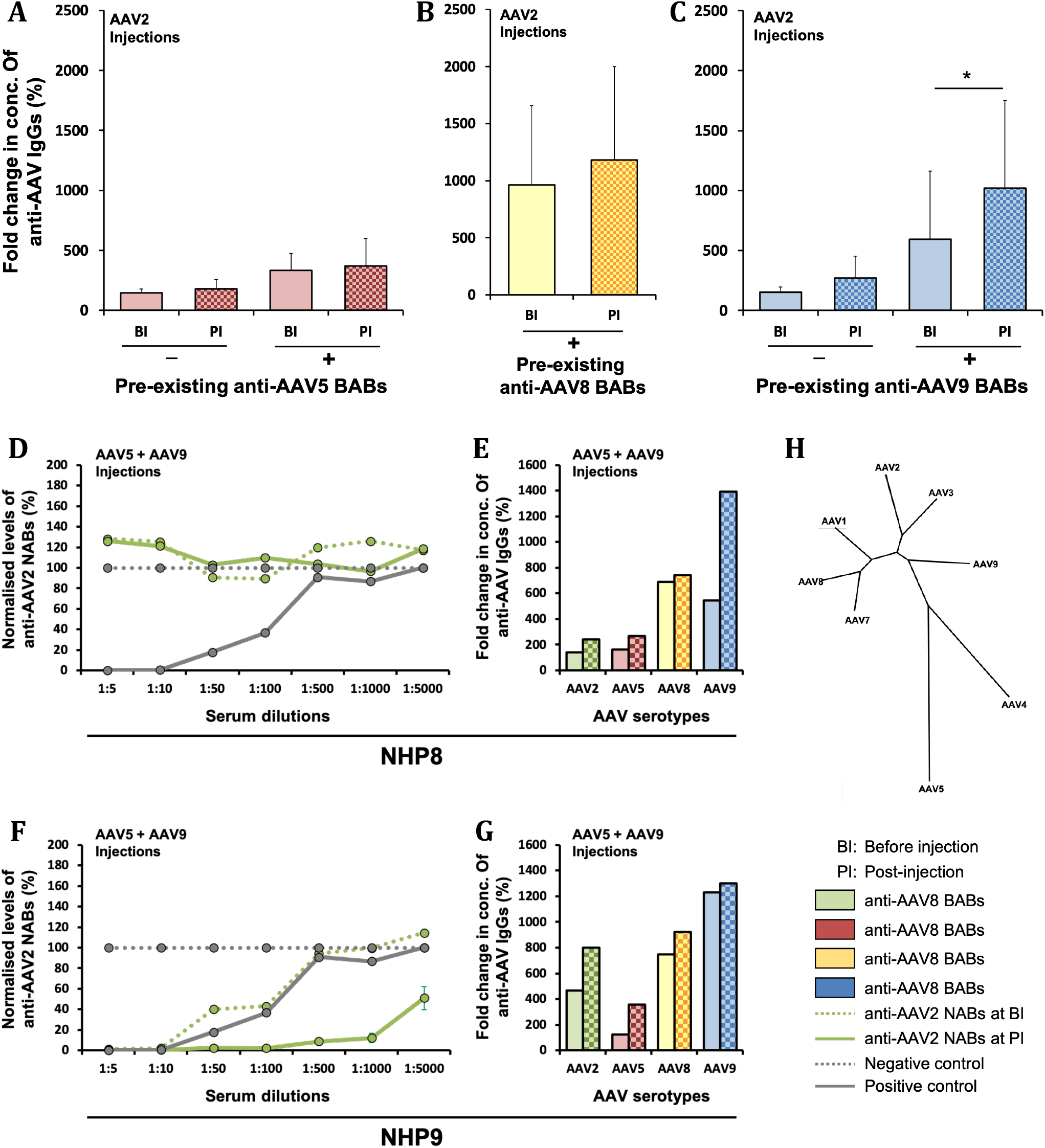
Cross-reactivity across serotypes. NHPs injected with AAV2 and tested before injection (BI) and post-injection (PI) for serum concentration of binding antibodies (A) against AAV5 in n=18 NHPs without (-) and n=9 NHPs with (+) pre-existing BABs; (B) anti-AAV8 in n=27 NHPs with (+) pre-existing BABs and (C) anti-AAV9 in n=7 NHPs without (-) and n=20 NHPs with (+) pre-existing BABs; (D,F) NABs against AAV2 and (E,G) BABs against AAV2, 5, 8 and 9 in (D,E) NHP8 and (F,G) NHP9 before and post-injection of AAV5+AAV9. Shown are mean values ±SD. Significance at individual time points was tested using Student’s t-test (*P< 0.05); (H) Phylogeny tree generated from AAV capsid sequences of different serotypes where the branch lengths are proportional to the evolutionary change (calculated using ClustalW).

### Influence of promoter and long-term monitoring of antibody levels

After we established the impact of dose on serum BAB and NAB levels, we investigated if there are other factors that can potentially have an effect. A group of animals that were injected with the same AAV serotype (AAV2) in each eye at the same dose (5.00E+11vg), were evaluated for their BAB levels post-injection. The only difference between these vectors was the promoter driving the transgene expression. 4 animals had a ubiquitous CAG promoter, whereas 2 animals had a ganglion cell specific promoter – SNCG^29^. 3 out of the 4 animals with the ubiquitous promoter had a higher level of BABs post-injection compared to the 2 animals with the ganglion cell specific promoter (Supplementary figure 3).

We monitored the same animals (Supplementary figure 3) long-term, by analyzing the BAB levels in serums collected at time-points of 1 month until 6 months post-injection. In 5 out of the 6 animals, the BAB levels increased 1M post-injection and this level stayed stable until 6M. 2 NHP samples that were tested at time points later than 6M (up to 1 year), also showed that the BABs could be detected until much later in the serums post-injection (data not shown).

There is an interest in re-administration of AAV^20,30^, hence lastly, we examined the effect of double and sequential injections on the post-injection antibody levels. In our cohort two animals had received double injections – NHP1 received an injection combining AAV2 along with AAV9 and NHP1 received sequential injections of AAV2 in the same eye. In both cases there was an increase in BABs post-injection, which stayed at the same level or increased only slightly after the second injection. This increase was comparable to two animals (NHP3 and NHP21) that had received similar serotypes and doses but as a single dose (Supplementary figure 2A).

## DISCUSSION

The pre-existing levels of anti-AAV antibodies in the serum have potentially important implications in study outcomes involving intraocular AAV administration. In this work we first examined the basal levels of anti-AAV antibodies against commonly used AAVs in the sera of macaques that have been used for pre-clinical testing of various gene therapy strategies. In humans anti-AAV2 NABs were found to be much higher than anti-AAV7 or 8^12,13^. AAV 4, 7, 8 and 9 originate from monkeys, whereas AAV2, 3 and 5 are believed to have a human origin^31,32^. Thus, it is not surprising that the levels of anti-AAV8 and anti-AAV9 antibodies were higher than the anti-AAV5 in the 41 NHPs we tested. As mentioned earlier the levels of AAV2 in our samples may have an exclusion bias, since very often animals in our studies are pre-selected to be AAV2 NAB negative prior to their enrollment to AAV based studies. Nonetheless, the anti-AAV5 levels in all samples tested were low, like what is observed in human samples. But, unlike humans who have low seroprevalence of AAV8 and AAV9, the anti-AAV8 and AAV9 levels were higher in NHPs^12^. Also, when the capsid sequence homology is compared, AAV5 is the most distantly related to all other serotypes (Figure6H). In our study we found that the most important factor that influences the rise in serum antibody levels is the dose injected in the retina (Figure 2), irrespective of the serotype (Supplementary figure 1) or the mode of injection (Figure 3). In the vitreal space there are higher chances of exposure to systemic circulation^15^, whereas the subretinal space does not expose AAV to systemic circulation owing to the blood-retina barrier^33,9^. Hence intravitreal AAV injections are expected to be more immunogenic than subretinal injections^34,35,18^. A study showed that IVT in one eye results in an immune response in the contralateral eye. This prevented a second IVT injection in the fellow eye, but a SR injection was possible^14^. On the other hand, a primate study concluded that subretinal injections of AAV8 in NHP induced both innate and adaptive immune responses^36^. In our study, we did not find a difference in the serum NAB and BAB levels in the animals that were injected by different modes of injection. There was an increase post-injection in both cases (Figure 3), but we only tested the antibody levels in the serum. It is possible that the differences (if any) may be more prominent locally than at a systemic level. Although, a study has shown a positive co-relation (R=0.3) between the NABs detected in the vitreal fluid and the NABs in the serum it did not make a comparison of BABs^26^. It may be worthwhile to test BABs in the vitreous fluid of NHPs to ascertain if there are indeed any differences between antibody levels after SR and IVT injections. This could pose a technical challenge though, as extracting the vitreal fluid is an invasive procedure that can affect the intraocular pressure, cause cataracts, or even result in retinal detachment^37^. Performing vitreous sampling before and after AAV injection, could result in more problems than the AAV injection itself.

An increase in dose refers to an increase in capsids, transgene and promoter. Each of these individually or collectively could be responsible for the increase in serum antibody titers^19,38,11^. We observed that response to capsid depends on the pre-existing levels of a particular serotype which differs by species. The response to transgene could not be evaluated as the samples were collected from different studies. But we were able to make a comparison of promoters in a group of macaques. Studies have shown that the promoter influences the AAV injection outcome, where ubiquitous promoters have been linked to higher toxicity affecting both the transgene expression and the function of the retina^21,22,19^. We observed a trend of higher serum antibodies in NHPs injected with ubiquitous promoters compared to cell-specific promoters (Supplementary figure 4).

The cross-reactivity between certain serotypes such as AAV2 and AAV9 could be because they are more closely related both in terms of genome sequence homology as well as the capsid sequence homology (Figure 6H). 84% genome sequence homology exists between AAV2 and AAV9, whereas the sequences of AAV2 and AAV5 have 54% homology, and AAV9 and AAV5 have a 46% homology^39,40^. This may have consequences in selecting the serotype that can be used in cases where there is a need for a second injection. For example, after a first injection with AAV2, a follow-up injection with AAV5 might entail less risks than a subsequent injection with AAV9. Our data, consistent with previous studies^26^ reveal that anti-AAV antibodies persist in the serum even after 6 months to 1-year post-injection, which has to be taken into consideration if repeat injections are required.

An immune response to viruses and generation of antibodies is a sign of a healthy functioning immune system, so the serum antibodies are not a cause for concern unless they cause an inflammation^21,11,41,42^. One of the important findings of our study was that in case of animals that developed severe local inflammation the serum antibodies were significantly higher than the animals that did not have any inflammation. This also correlated with clinical signs of inflammation (Figure 3), and with the transgene expression (Table1) Hence, the serum antibody levels can be used as a relatively less invasive indicator for ocular inflammation.

Owing to ethical issues, high costs, and inter-individual variability it is challenging to achieve statistical significance in studies involving non-human primates. Nonetheless, with our comprehensive retrospective analysis of 41 NHPs used in preclinical gene therapy studies over a period of 10 years, we were able to make some statistically significant observations, and show some trends which, when considered in the context of other studies provide meaningful insights for future pre-clinical studies involving AAV-mediated gene delivery.

## ACKNOWLEDGEMENTS

This work was supported by ERC Starting Grant (REGENETHER 639888), the Institut National de la Santé et de la Recherche Médicale (INSERM), Sorbonne Université, The Foundation Fighting Blindness, Agence National de Recherche (ANR) RHU Light4Deaf, *LabEx LIFESENSES (ANR-10-LABX-65) and *IHU FOReSIGHT (ANR-18-IAHU-01). The authors would like to thank Melissa Desrosiers and Camille Robert for their technical assistance with production of plasmids and viral vectors. The authors also extend their thanks to Shivani Shah and Oudom Kem for writing and implementing the code in R.

## CONFLICT OF INTEREST

DD is an inventor on a patent of adeno-associated virus virions with variant capsid and methods of use thereof with royalties paid to Adverum Biotechnologies (WO2012145601 A2), patent applications on noninvasive methods to target cone photoreceptors (EP17306429.6 and EP17306430.4) licensed to Gamut Tx (now SparingVision). DD is a founder of Gamut Tx (now SparingVision). The remaining authors declare that the research was conducted in the absence of any commercial or financial relationships that could be construed as a potential conflict of interest.

## MATERIAL AND METHODS

### AAV Production

AAV vectors containing transgenes were packaged by co-transfection in HEK293 cells (ATCC CRL-1573), harvested 3 days post transfection cells and purified by iodixanol gradient ultracentrifugation. The 40% iodixanol fraction was collected after a 90 min spin at 354000 g. Concentration and buffer exchange were performed against PBS containing 0.001% Pluronic^32^. AAV vector stocks titers were then determined based on real-time quantitative PCR titration method using ITR primers and SYBR Green (Thermo Fischer Scientific) as described earlier^43^.

### Animals and Intraocular injections

All non-human primates in this study were Cynomolgus *Macaca fascicularis* and originated from Mauritius. Prior to injections, they were anesthetized with an intramuscular injection of Ketamine, 10mg/kg (Imalgene 1000, Merial) and Xylazine 0.5mg/kg (Rompun 2%, Bayer). Anesthesia was maintained with an intravenous injection of propofol at 1ml/kg/h (PropoVet Multidose 10mg/ml, Zoetis). Pupils were dilated using 1% tropicamide (Mydriaticum, Théa Pharmaceauticals, France) and the eyelids were kept open using eyelid speculums. A 1-ml syringe equipped with a 32-mm, 27-gauge needle was used for intravitreal injections, by insertion into the sclera approximately 2 mm posterior to the limbus to deliver between 50-100 μl of the viral vector solution. For subretinal AAV injections, two 25-gauge vitrectomy ports were set approximately 2 mm posterior to the limbus, one for the endo-illumination probe and the other for the subretinal cannula. A 1-ml Hamilton syringe equipped with a 25-gauge subretinal cannula with a 41-gauge tip was used for the injection. The endoillumination probe and cannula were introduced into the eye. 50-100 μl of the viral vector solution was injected subretinally to create a bleb either below or above the fovea. After subretinal or intravitreal vector administration, opthtalmic steroid and antibiotic ointments (Fradexam, TVM) were applied after injections. All animal experiments and procedures were ethically approved by the French “Ministère de l’Education, de l’Enseignement Supérieur et de la Recherche” and were carried out according to institutional guidelines in adherence with the National Institutes of Health guide for the care and use of laboratory animals as well as the Directive 2010/63/EU of the European Parliament.

### Serum collection and dilutions

NHP blood was collected from a peripheral vein using a 22G into a red top vacutainer, at various time points before and after ocular injections of AAVs. The blood sample was centrifuged at 800g for 10mins, and the serum (supernatant) was transferred into a separate Eppendorf tube and stored at -80°C until further use. The serum was diluted in blocking buffer (6% milk in 1xPBS) for ELISA and in cell culture medium (DMEM+10% Fetal Bovine Serum) for NAB Assays.

### NAB Assay

NAB Assay for AAV2 was performed using HEK293T cells (ATCC CRL-1573) and for AAV9 using LEC2 cells (ATCC CRL-1736) as described before^24^. Briefly, HEK293T cells were plated at a density 7.00E+5/well and Lec2 cells were plated at a density of 6.50E+5/well in a 96 well plate and placed in an incubator (37°C/5% CO_2_) for 4h. In another plate serum dilutions in DMEM+10% FBS were prepared. To these serum dilutions AAV2 at a multiplicity of infection (MOI) of 6400 and AAV9 at a MOI of 20000 was added and incubated for 2h at 4°C. The serum-virus mix was added to the cells in duplicates and incubated overnight at 37°C. 24h later the cells were lysed using Cell culture lysis buffer (Promega E1531) and mixed with luciferase assay reagent (Promega E1501) as per the manufacturers protocol. The luminescence was measured using the Spark Multimode microplate reader (TECAN, Switzerland).

### ELISA

A 96-well plate (Nunc Maxisorp, ThermoFischer scientific 442404-21) was coated with 1.00E+9 vg AAV/well. To generate a standard curve a dilution series using the Monkey IgG-UNLB antibody (Southern Biotech, 0135-01) was coated on each plate and incubated overnight at 4°C. The AAV and antibody was diluted in coating buffer (0.1 M carbonate buffer pH 9.5). The plate was washed with blocking buffer (6% milk in 1xPBS), followed by incubation with serum in triplicates for 2h, washing with wash buffer (1xPBS+0.05% Tween) and incubation with secondary antibody - Mouse-anti-monkey IgG-HRP (Southern Biotech, 4700-05) for 1h. For visualization Tetramethylbenzidine (TMB) substrate (Sigma Aldrich, T0440) was applied for 10mins, the reaction was stopped using the TMB stop solution (ThermoFischer Scientific, SS04) and the luminescence measured at 450nm using the Spark Multimode microplate reader (TECAN, Switzerland).

### Statistical analysis

Statistical analysis was performed using Prism 9.0 software (GraphPad, San Diego, CA, USA). All data are presented as mean ± standard deviation (SD). The number of samples (N) used for individual experiments is given in the figure legends. Student’s t-tests were used to test the significance between means of groups. A *P*-value below 0.05 was considered significant and indicated on graphs as *P< 0.05, ** P< 0.01, ***P< 0.001, **** P< 0.0001. In case of NAB assays, to estimate the serum dilution at which 50% neutralization occurs, we fit a curve between observations using linear interpolation. This computation for interpolation has been performed using the ‘*stats*’ package inbuilt in R^44^.

## FIGURE LEGENDS

**Figure S1.**
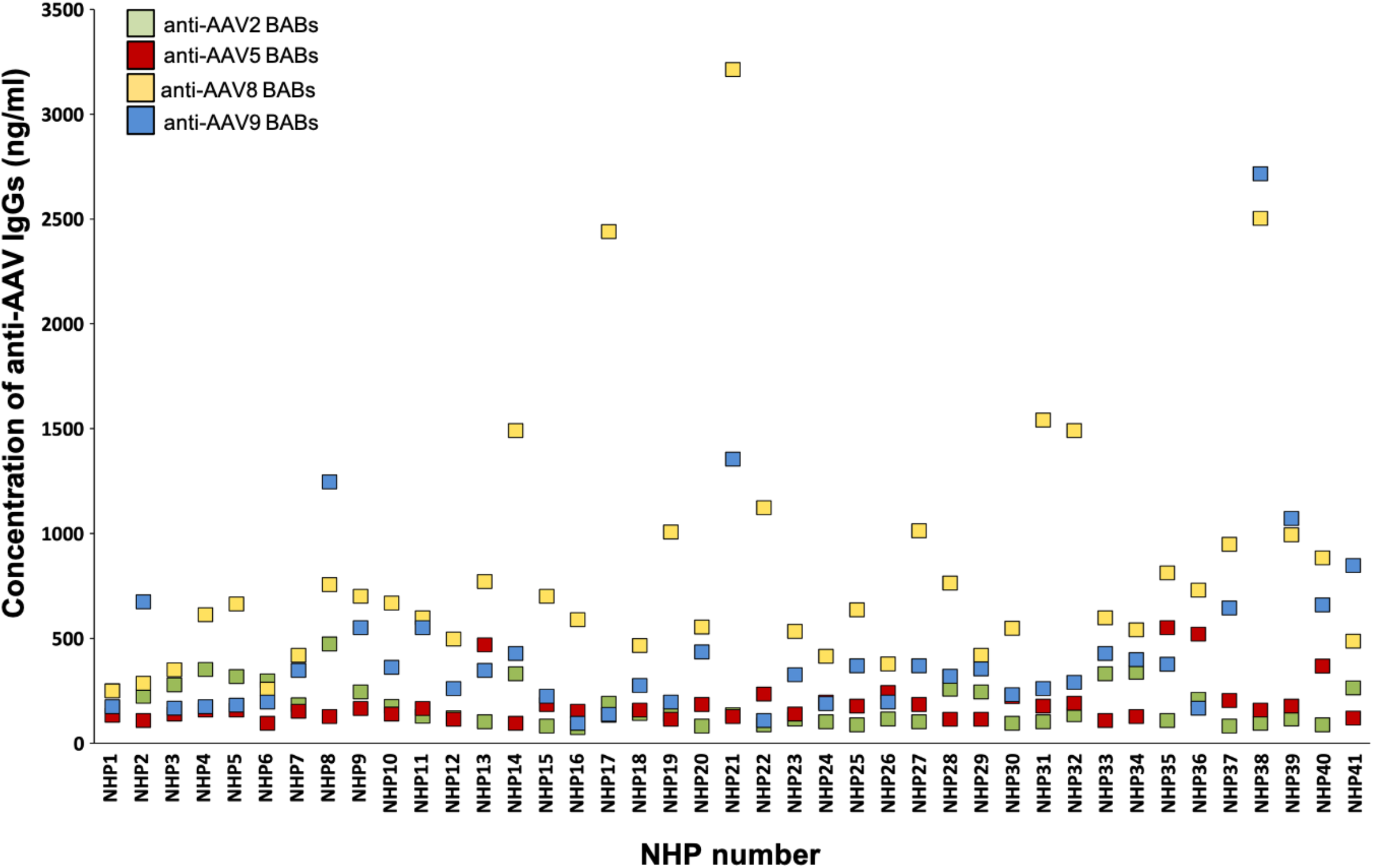
Higher basal levels of BABs against AAV8 and AAV9 in NHP serums. Concentration of BABs against AAV2 (green), AAV5 (red), AAV8 (yellow) and AAV9 (blue) in serums from 41 NHPs before injection shown individually.

**Figure S2.**
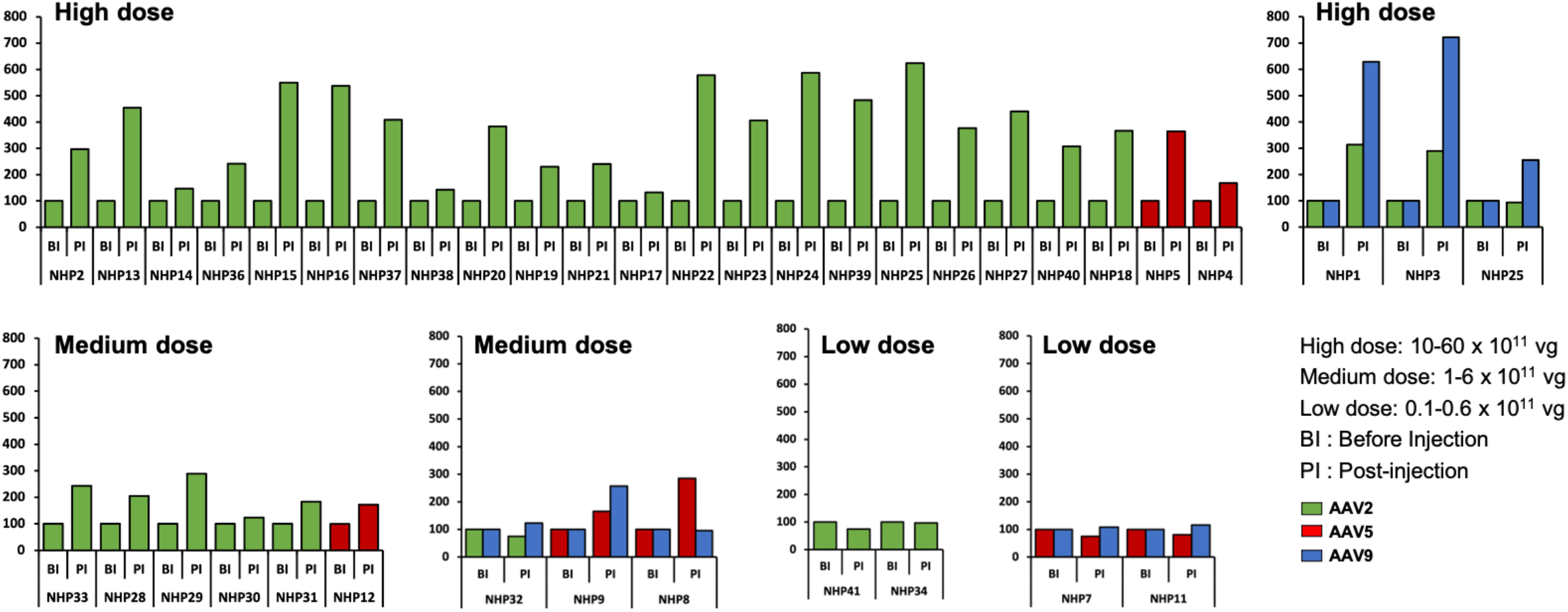
Dose dependent increase in levels of BABs post-injection. irrespective of the AAV serotype injected. Fold change in the concentration of BABs against AAV post injection (PI) shown for each animal individually. The BAB levels are measured for the same serotype that was injected (indicated by the color of the bar, AAV2: green, AAV5: red and AAV9: blue) at high, medium or low dose (as indicated). The values for BABs are normalized relative to the level before injection (BI), which is set to 100. High dose: 10-60 × 10^11^ vg, Medium dose: 1-6 × 10^11^ vg, Low dose: 0.1-0.6 × 10^11^ vg.

**Figure S3.**
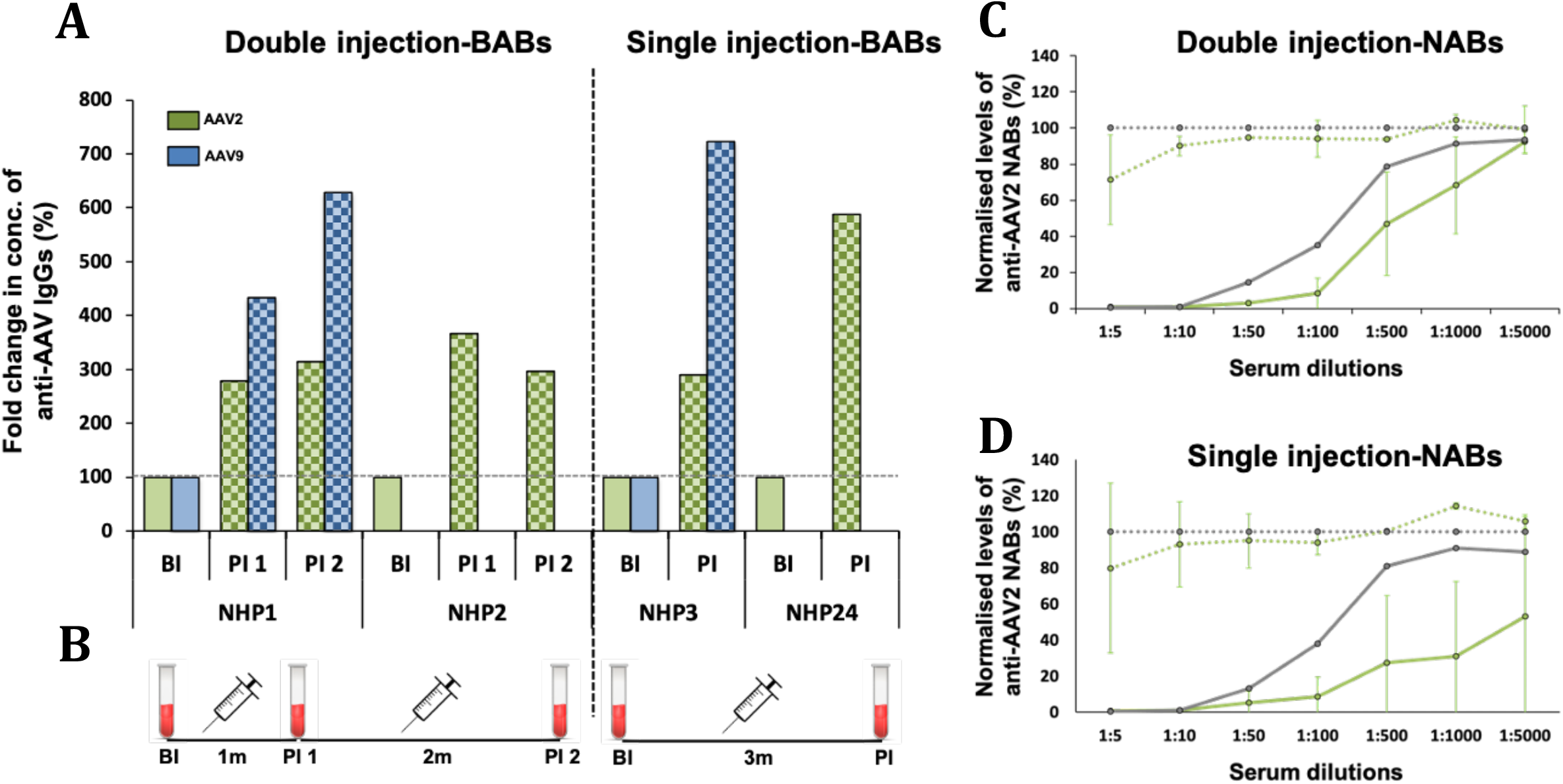
Effect of re-administration of AAV injection on BAB and NAB levels. (A) Change in the concentration of BABs against AAV2 (green bars) and AAV9 (blue bars) in NHP1 (injected with AAV2+AAV9), NHP2 (injected with AAV2), NHP3 (injected with AAV9+AAV2) and NHP24 (injected with AAV2); (B) Schematic representation of serum collection points for double and single injections, BI: before injection, PI: post injection, PI1 and PI2: post-injection 1 and 2 respectively; Anti-AAV2 NAB levels in (C) NHP1 and NHP2 that received double injections and in (D) NHP3 and NHP24 that received a single injection.. The values for BABs are normalized relative to the level at BI (set to 100). The values for NABs are normalized relative to the negative control (set to 100), and shown as Mean ±SD. All 4 animals received a high dose with a total dose ranging between 1.40-3.00 E+12 vg.

**Figure S4.**
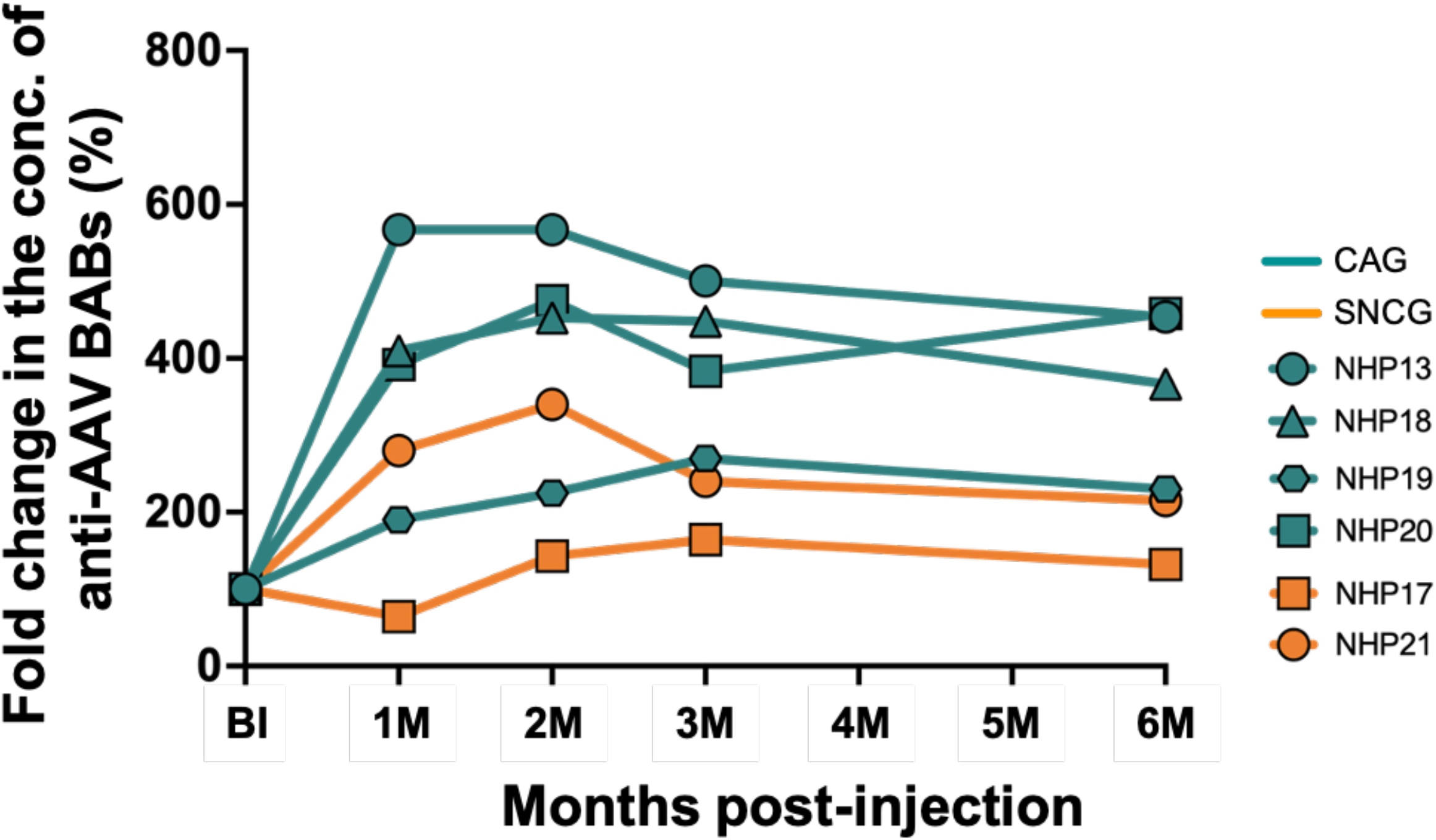
Increase of BAB production in NHPs injected with transgene containing an ubiquitous promoter. Change in the concentration of BABs against AAV2 in 3 NHPs injected with a transgene containing a ubiquitous promoter (CAG) and 2 NHPs injected with a transgene containing a ganglion cell-specific promoter (SNCG). All animals received the same serotype (AAV2-7m8) and viral dose in each eye (5·00E+11 vg in each eye) as a single injection. The values for BABs are normalized relative to the level before injection (BI), which is set to 100.

